# Lifelong learning of cognitive strategies for physical problem-solving: the effect of embodied experience

**DOI:** 10.1101/2021.07.08.451333

**Authors:** Kelsey R. Allen, Kevin A. Smith, Laura-Ashleigh Bird, Joshua B. Tenenbaum, Tamar R. Makin, Dorothy Cowie

## Abstract

‘Embodied cognition’ suggests that our bodily experiences broadly shape our cognitive capabilities. We study how embodied experience affects the abstract physical problem-solving strategies people use in a virtual task where embodiment does not affect action capabilities. We compare how groups with different embodied experience – 25 children and 35 adults with congenital limb differences versus 45 children and 40 adults born with two hands – perform this task, and find that while there is no difference in overall competence, the groups use different cognitive strategies to find solutions. People born with limb differences think more before acting, but take fewer attempts to reach solutions. Conversely, development affects the particular actions children use, as well as their persistence with their current strategy. Our findings suggest that while development alters action choices and persistence, differences in embodied experience drive strategic changes in the acquisition of cognitive strategies for balancing acting with thinking.

**Statement of Relevance:** Theories of embodied cognition suggest our cognitive and perceptual capabilities are shaped by how our bodies constrain our interactions with the world; however, these tasks often study short-term effects or setups where body differences impact task solutions. Here we compare the performance of children and adults with and without congenital limb differences (missing hands/upper limbs) on a virtual physical problem-solving task, where both groups had equal motor capabilities for interacting with the world. Across ages, participants with limb differences solved these tasks as proficiently as those without limb differences, but took fewer attempts to come to solutions, with more time spent thinking between attempts. This suggests that early life differences in embodied experience cause changes in individuals’ “cognitive strategies” for allocating time between thinking and acting.

## Introduction

Everyday experience is both constrained and enabled by the bodies we inhabit. Taller people can reach further, while people with two fully functioning hands can manipulate multiple objects at the same time. ‘Embodied cognition’ (Wilson, 2002) suggests that such constraints play a fundamental role in broadly shaping our cognitive and perceptual experiences. Many versions of embodiment theory suggest that these effects reach beyond the types of experiences we have into how we reason about those experiences. Supporting this view, researchers have shown that when individuals’ bodies or skills are altered, e.g. through temporary training or by being born with limb differences, this can change their perceptual capacities (Aglioti, Cesari, Romani, & Urgesi, 2008; Hagura, Haggard, & Diedrichsen, 2017), spatial cognition (Makin, Wilf, Schwartz, & Zohary, 2010), body representation (Maimon-Mor, Schone, Moran, Brugger, & Makin, 2020), or motor skills (Maimon-Mor, Schone, Slater, Faisal, & Makin, 2021). Here we ask if these effects of embodiment can be broader, by testing whether differences in embodied experience (through limb differences) affect the ways that people think about acting in the world, even when their capacities for action are made equal.

Prior studies have rarely addressed how a lifetime of embodied experience affects a general choice of “cognitive strategies”: how people allocate cognitive resources to thinking about and acting in the world. Through short-term direct manipulations of people’s bodies or accessible actions, researchers have shown that people are sensitive to action costs – the amount of effort required to perform actions – for both motor planning (Izawa, Rane, Donchin, & Shadmehr, 2008) and motor behaviors (Prévost, Pessiglione, Météreau, Cléry-Melin, & Dreher, 2010). Priming people with different action costs in a perceptual decision-making task can even affect their decisions in a setting where action costs no longer apply (Hagura et al., 2017), suggesting that action costs can generalize beyond the immediate task. However, these studies transpire on the scale of minutes, and typically demonstrate behavioral effects where costs are manipulated within a task, but do not show that long-term and generalized action costs are learned from embodied experience.

To study the effect of embodied experience over longer time-scales, researchers have investigated the perceptual and motor capabilities of individuals born with limb differences. However, the tasks used to study these capabilities often require judgments related to absent body parts, and therefore differences in behavior might be driven by differences in sensorimotor experience or available information. For example, while people with congenital limb differences are slower to judge whether a picture is of a left or right hand (Maimon-Mor et al., 2020; though c.f. Vannuscorps & Caramazza, 2016; Vannuscorps, Pillon, & Andres, 2012), they lack first-person experience of that hand.

Here we test the hypothesis that the subtleties of growing up with a different body may affect the everyday cognitive strategies people use to solve problems in their environments, even when the capabilities tested are divorced from particular bodily differences. For example, if people with congenital limb differences have learned that actions are in general more costly – because, perhaps, it is more difficult to use artifacts designed for people with two hands (see Figure 1) – we might expect that they will differ in how they approach physical problems generally, even when action costs are equated. Such cognitive strategies for action have been observed previously on shorter time-scales: e.g., individuals adapt their motor plans to their own levels of uncertainty and variability (Harris & Wolpert, 1998; Gallivan, Chapman, Wolpert, & Flanagan, 2018; Körding & Wolpert, 2004), including becoming more persistent (Leonard, Lee, & Schulz, 2017), or spending more time thinking before acting (Dasgupta, Smith, Schulz, Tenenbaum, & Gershman, 2018). The differences in cognitive strategies might be expected to emerge early in infancy, when experience begins driving motor skill acquisition (Adolph, Hoch, & Cole, 2018), but might also develop throughout childhood alongside more precise motor planning, control and tool use (Berard & Vallis, 2006; Chicoine, Lassonde, & Proteau, 1992; Adalbjornsson, Fischman, & Rudisill, 2008).

**Figure 1.**
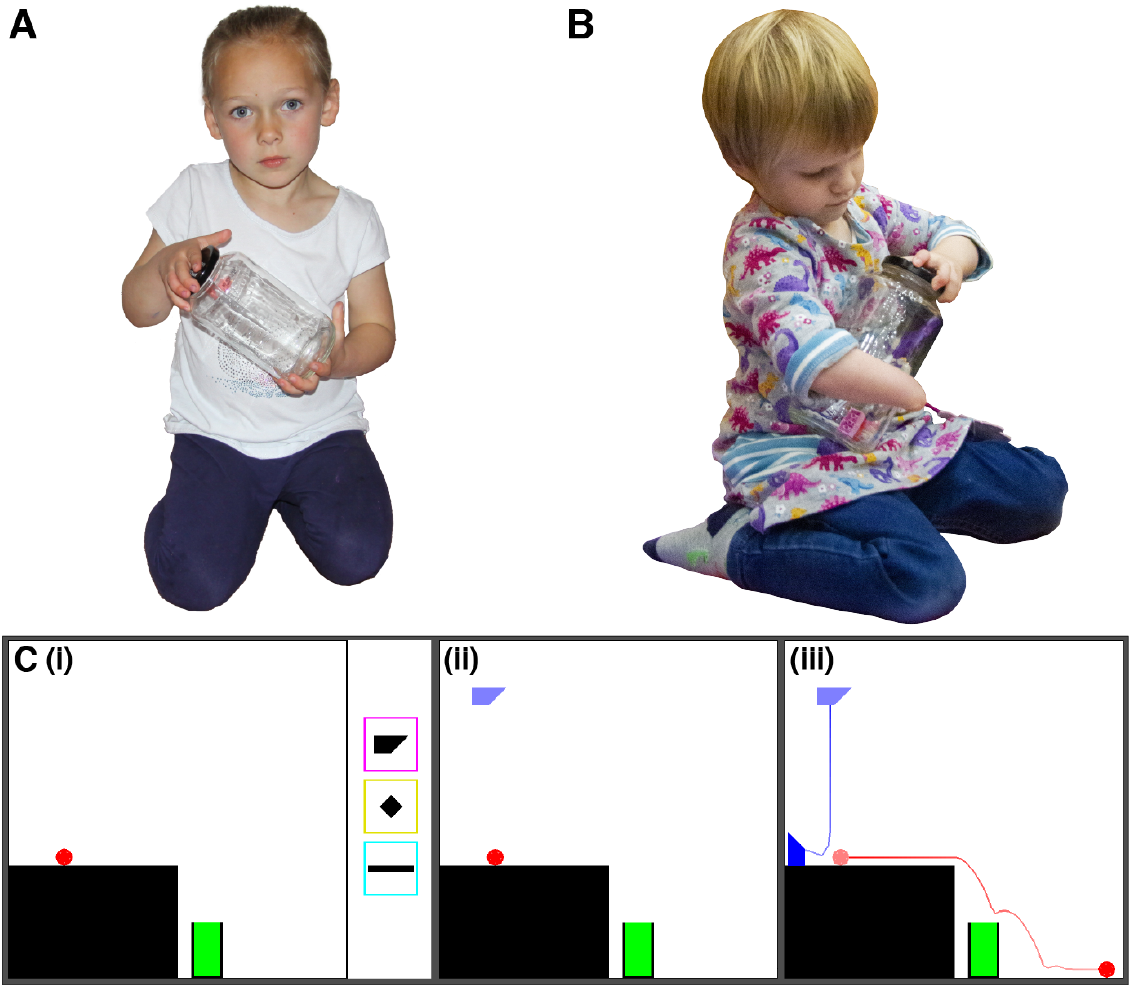
Individuals with limb differences must engage in independent motor problem-solving for many everyday behaviours, like opening a jar. These experiences may change their cognitive strategies for motor tasks in general. For example, with two hands, opening a jar can be accomplished by using one hand to stabilize the jar while the other one twists the lid (**A**). With a single hand (**B**), opening a jar can be accomplished by using one’s arm and torso to stabilize the jar. The Virtual Tools game (**C**) equalizes action possibilities and costs for individuals with different types of limbs by creating a virtual action space. (i) The aim is to move the red ball into the green goal. A participant selects a tool from three options (shapes in coloured boxes) and places it in the scene (ii). Once placed, physics is “turned on” and objects can fall under gravity or collide with each other (iii); the blue and red lines represent the observed motion trajectories for the tool and the ball respectively.

To test the influence of embodied experience on cognitive strategy learning, we studied behavior in a *virtual* physical problem-solving task where all participants had equal capabilities to interact with the world. This “Virtual Tools game” (Allen et al., 2020) requires people to use virtual objects as tools to solve a physical problem (e.g., getting the red ball into the green goal, Figure 1C) using a single limb to control a cursor, thus equating action costs. Allen et al. (2020) provide a set of performance metrics that describe how people solve these puzzles to measure the cognitive strategies recruited for the task.

We chose participant groups to represent a diverse range of embodied experience: children (5-10-year-olds) and adults born with limb differences, and age-matched children and adults born with two hands. Children have less embodied experience than adults, while individuals with limb differences have dramatically different kinds of experience. By using a virtual task with simple controls, we equate manipulation capabilities and instead study how embodied experience affects the cognitive strategies that support planning and reasoning for action more generally.

We tested whether cognitive strategies are affected by life experience, as indexed by age (children versus adults) as well as limb differences. We first predicted that those having to devise unique solutions to everyday physical problems due to growing up with a limb difference might use different, and perhaps even more efficient, strategies while solving the virtual puzzles. To assess this, we considered the key outcome measures of action type, thinking time, attempts to solution, time to solution, and solution rate introduced by Allen et al. (2020). Second, given that tool use capabilities develop from early to late childhood (Beck, Apperly, Chappell, Guthrie, & Cutting, 2011; Keen, 2011), we expected solution rates to improve from children to adults. Finally, we tested whether differences in cognitive strategies between those with and without limb differences would emerge early because compensatory behavior evolves early, or whether differences might grow with development as motor and cognitive skills develop. Overall, we found that participants with limb differences do use a different set of cognitive strategies than those without – spending more time thinking while interacting less with the world. While performance does improve with age, we did not find evidence that the thinking/acting difference between participants with and without limb differences changes over development.

### Open Practices Statement

This study was not preregistered. The data and analyses have been made available on a permanent third-party archive https://github.com/k-r-allen/embodied_experience. The stimuli for the study can be found at https://sites.google.com/view/virtualtoolsgame and in the Supplemental Materials.

## Methods

### Participants

We recruited a total of 145 participants across four groups: 40 adults without limb differences (Adult-NLD), 35 adults with limb differences (Adult-LD), 45 children with no limb differences (Child-NLD), and 25 children with limb differences (Child-LD). LD and NLD participants were well matched for age (Child-LD mean: 7.91yo, sd: 1.84; Child-NLD mean: 7.94yo, sd: 1.74; Adult-LD mean: 40.7yo, sd: 15.5; Adult-NLD mean: 41.2yo, sd: 15.2). We extensively liaised with two UK charities for supporting children with a limb difference and their families, as well as using our ties with the limb-difference community and existing volunteer databases to recruit as many children and adults with congenital upper limb differences as possible over the research period. We included participants with congenital upper limb anomalies, as summarized in Tables S2 and S3 in the Supplemental Material. Information on limb differences was self-reported, but verified for a subset of participants (88% and 55% in children and adults, respectively). We recruited children and adults without limb differences to match the education level and age of the special population over the same period. We recruited a larger sample of children without limb differences to give us a greater understanding of the range of typical performance in a two-handed population. Two-handed children were recruited through an existing university volunteer families database and affiliated Facebook page. For all participants, participation was approved by university ethics boards.

**Table 1.** Proportion of children and adults with limb differences who have an affected right or left arm, or both arms, and associated ages of groups in years.

| Age Group | Affected Limb(s) |  |  | Mean(SD) Age |
| --- | --- | --- | --- | --- |
|  | Right Arm | Left Arm | Both |  |
| Children | 0.32 | 0.44 | 0.24 | 7.91 (1.84) |
| Adults | 0.42 | 0.55 | 0.03 | 40.7 (15.5) |

### Experiment

As with Allen et al. (2020), the experiment was run online on participants’ personal computers at home. All participants were provided with an identifying code used to link their performance with individual information (e.g., specific limb differences). The experiment progressed through two stages: motor pre-test, and Virtual Tools game. We also collected several additional demographic and clinical details (see below).

All participants were given the same experiment, with only three exceptions that differed between children and adults: (1) children received simplified instructions for all stages, (2) adults played one additional Virtual Tools level that we removed from the children’s experiment due to excessive challenge (see Section S3 in the Supplemental Material), and (3) adults were given a more extensive questionnaire that included additional questions about the strategies they had used and video games they had played before.

#### Motor pre-test

The motor pre-test was used to measure participants’ ability to control the cursor. Each trial began with a star centered in a 600×600px area on the screen. Once the star was clicked, a 10px radius circle appeared in a random position either 150px or 250px from the center. Participants were instructed to click on the circle as quickly and accurately as possible. Participants completed 10 motor test trials (five at each distance from the center).

On each trial we measured (a) reaction time, and (b) the distance (in px) between the center of the circle and the cursor click location. As a measure of participants’ basic motor accuracy, we took the median of both of those measures across all 10 trials; we used the median to avoid skew from outlier trials, and found in pilot testing that this was a relatively stable measurement.

#### Virtual Tools game

On each level of the Virtual Tools game, participants were presented with a scene and three “tools” (see Figure 1C-i), along with a goal condition (e.g., “get the red object into the green goal area”). Participants could accomplish this goal by clicking on a tool and then an unobstructed part of the game area to place that tool (Figure 1C-ii); as soon as the tool was placed, physics was “turned on” and objects could fall under gravity or collide with each other depending on the specific tool placement made (Figure 1C-iii). If the goal was not accomplished, participants could press a button to reset the scene to its initial state. Participants could attempt to solve the level as many times as they liked, but were limited to a single tool placement for each attempt. Participants could move onto the next level once they had accomplished the goal, or after 60 seconds had passed.

Following Allen et al. (2020), participants were initially given instructions about how the game functioned, including the difference between static (black) and moving (blue/red) objects, goal areas, and how to place tools and reset the level. For familiarization with the interface and physics, participants were given one introductory level that required them to place tools at least three times without a goal, followed by two simple levels that they were required to solve but that were not analyzed. For adults, this process was identical to that of Allen et al. (2020); children received simplified instructions that conveyed the same information (see Figure S1 in the Supplemental Material).

In the main task, participants were asked to solve 14 different levels, each with a different set of three tools, designed to probe knowledge of diverse physical principles (e.g., support, collisions, tipping; see Figure 2). As in Allen et al. (2020), on each level, we recorded all attempts from each participant – defined as the tool chosen and where it was placed; the time elapsed between the start of the level and the time of the attempt; and whether the level was solved.

**Figure 2.**
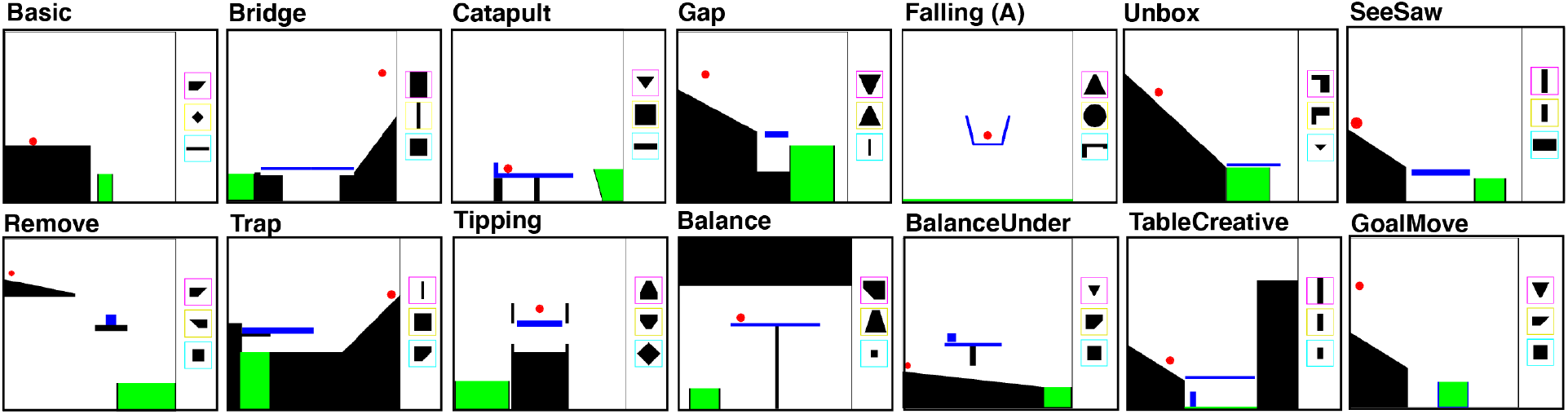
The fourteen levels of the Virtual Tools game (Allen, Smith, & Tenenbaum, 2020) that participants played. These cover a wide variety of physical action concepts including “balancing,” “launching,” “catapulting,” “supporting,” and “tipping.” To play the game, please see https://sites.google.com/view/virtualtoolsgame.

For the main analyses, we used four different overall performance metrics defined by Allen et al. (2020) to measure different facets of performance: (1) whether the level was solved (solution rate), (2) time until the solving attempt was performed (time to solution), (3) how many attempts were taken until the level was solved (attempts to solution), (4) times to the first attempt and the average time between attempts (thinking time). We also analyzed the specific kinds of attempts taken (attempt type) using the same methodology introduced in Allen et al. (2020) to cluster and classify participants. All measures were automatically extracted from the recorded tool placements and timings.

#### Additional measures

After both the motor test and Virtual Tools task, participants were given a short questionnaire to ask what device they used to control the cursor, and, for participants with limb differences, how the cursor was controlled. Additionally, for the adult participants we included the questions asked in Allen et al. (2020), including prior video game experience, and free-form responses about strategies they had used on the task. In separate surveys, we gathered demographics and interface information (Section S2.2 in the Supplemental Material), limb-differences information (Section S2.3 and Tables S2 and S3), and verbal and nonverbal IQ (for a subset of children only; Section S2.1). Age was used as a covariate in our analyses, while gender, device, and limb usage were studied as possible moderators (see Section S7 in the Supplemental Material). Free-form strategy descriptions were read in order to discover non-standard ways of solving puzzles, but were not directly analyzed.

### Analysis

We analyzed the motor test performance using linear models to predict the dependent variable (either median reaction time or click distance) as a function of both age group and limb difference group, controlling for the effect of differences due to age in years separately for adults and children.

For all of the Virtual Tools performance metrics, variables were modeled using linear mixed effect models, using random intercepts for participants and levels. Additionally, we included age in years and median motor test response times as covariates, parameterized separately for children and adults. We had prespecified these as covariates since we believed that they would account for general performance differences; nonetheless analyses without covariates produced similar results (see Section S6 in the Supplemental Material). For all but the solution metric, we conditioned our analyses only on *successful* levels, as we were interested in the mental processes that led to solutions, and not processes that might be indicative of frustration or perseverance; however, analyzing all levels produced a qualitatively similar pattern of results (see Section S8 in the Supplemental Material).

In some analyses we attempt to differentiate the *type* of participants’ attempts, either to test whether the type of attempt is different across groups, or to test whether participants are switching types between attempts. In order to classify attempts in a data-driven way, we use the classification methodology of (Allen et al., 2020) that groups attempts using nonparametric clustering. To compare attempt types across groups, we applied a leave-one-out classification analysis where, for each participant, we formed probability distributions over the tool identities and spatial positions by all other members of their group and those from members of the other group, then calculated the relative likelihood that the tool placement for a given level was a member of the correct group. To investigate whether attempt types change from attempt to attempt, we formed clusters over tool spatial positions using all attempts from all participants within a level. Two consecutive attempts were considered a “switch” if each attempt came from a separate cluster.

## Results

### Basic motor abilities across groups

We used the motor pre-test to examine whether there were any group differences in cursor control which could affect performance on the Virtual Tools game. Children could control the cursor, with an average pixel error of 7.65*px* (95% CI=[6.74, 8.55]) and reaction time of 3.04*s* (95% CI=[2.67, 3.41]), albeit less accurately and more slowly than adults (error: 2.92*px*, 95% CI=[2.52, 3.31]; RT: 1.91*s*, 95% CI=[1.73, 2.09]). Exploratory analysis on the linear models showed that when treating age as a continuous variable, children’s control improved linearly with age (*t*(65) = 3.09, *p* = 0.003), reaching adult-like levels by 9-10 years old (see Figure 3).

**Figure 3.**
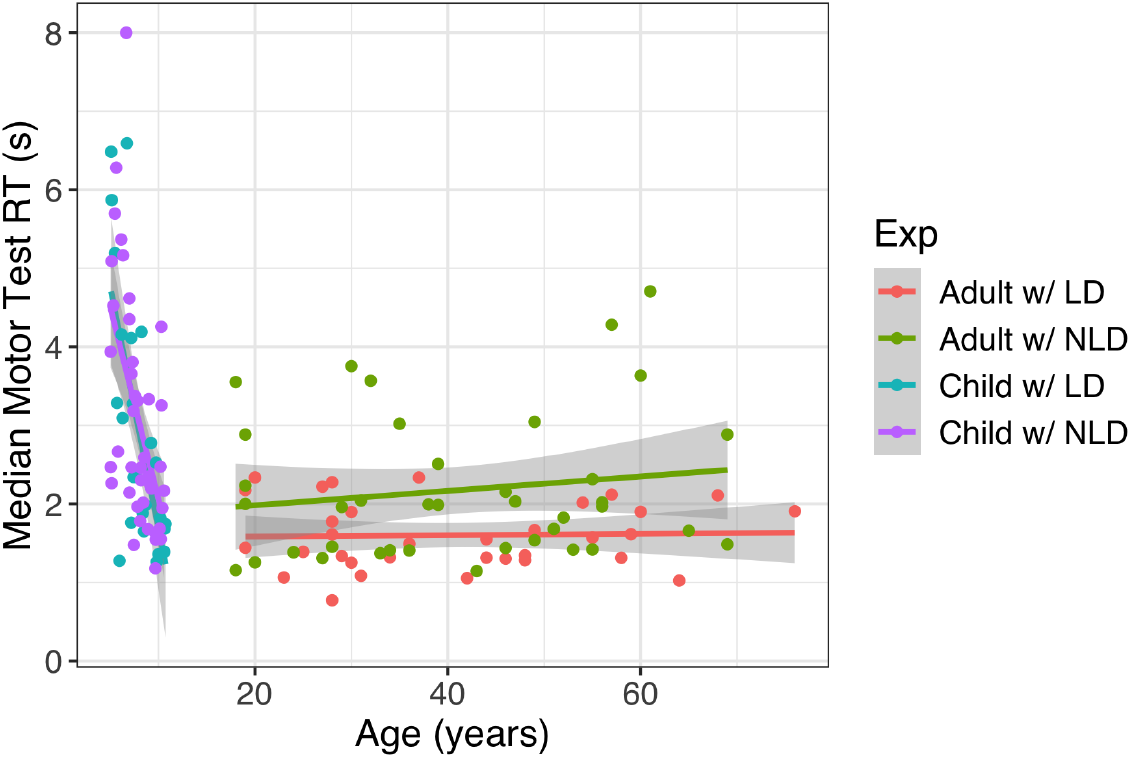
Reaction time on the motor task by participant age and group. Participants with limb differences were slightly *faster* on this task, suggesting that any differences in time to act or solve problems in the Virtual Tools game are not driven by differences in cursor control capabilities.

Importantly, individuals with and without limb differences performed comparably on both motor error and reaction time. While there was a difference in median click-time between participants with and without limb differences, participants with limb differences were slightly *faster* (by 356*ms*, 95% CI=[14, 698]; *F*(1, 135) = 4.15, *p* = 0.044), though they clicked marginally further away from the target (by 0.79*px*, 95% CI=[-0.14, 1.72]; *F*(1, 135) = 2.78, *p* = 0.098). There was no interaction found between age group and limb difference for either motor speed (*F*(1, 134) = 1.76, *p* = 0.19) or error (*F*(1, 134) = 0.001, *p* = 0.98). Note that the differences found in motor control were relatively inconsequential for the Virtual Tools game – 0.79*px* additional error would have little effect on 600 × 600*px* game screens, and an extra 356*ms* would be hard to detect with an average time between attempts of over 10*s*. We therefore showed that both groups should have a level playing field for interacting with the Virtual Tools game.

### Age, but not limb difference, impacts overall performance across levels in the Virtual Tools game

Moving onto the Virtual Tools game, we initially tested whether limb difference and age affected overall solution rates or time (in seconds) to reach a solution. We found gross differences in solution rates between children and adults (adults: 85%, children: 77%; *χ*^2^(1) = 11.2, *p =* 0.0008; Figure 4), but no effect of limb difference (*χ*^2^(1) = 1.21, *p =* 0.27), nor any interaction between age and limb group (*χ*^2^(1) = 0, *p* = 0.99). Exploratory analysis showed that treating age as a continuous variable additionally predicted success in a way that differed between children and adults (*χ*^2^(2) = 12, *p* = 0.0025), with children’s solution rates improving with age (log-odds increase per year: 0.290, 95% CI = [0.038, 0.543]), and adults’ performance getting worse (log-odds decrease per year: 0.029, 95% CI = [0.006, 0.053]).

**Figure 4.**
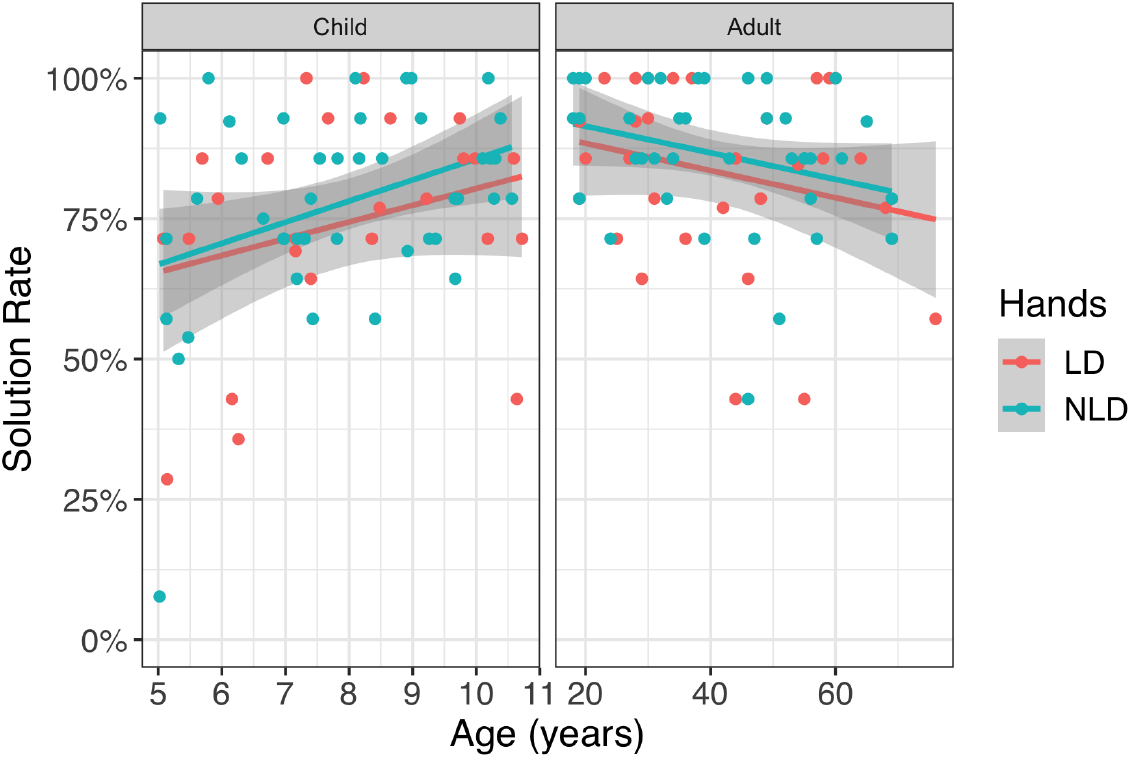
Solution rate (percentage of levels solved by each participant) as a function of age for children (left) and adults (right) with limb differences (LD) and with no limb differences (NLD). Grey areas represent standard error regions on the regression lines. Children’s solution rate improves as they age (log-odds increase per year: 0.290, 95% CI = [0.038, 0.543]), and adults’ performance worsens (log-odds decrease per year: 0.029, 95% CI = [0.006, 0.053]). There is no effect of limb difference on solution rates (*χ*^2^(1) = 1.21, *p* = 0.27).

The time to solve these levels shows a similar pattern of results: overall, children were slower than adults to find a solution (*χ*^2^(1) = 40.0, *p* = 2.6 * 10^−10^; Figure 5B), but we did not find evidence that limb differences affect the time to solve the levels (*χ*^2^(1) = 0.41, *p* = 0.52), nor was there an interaction between limb group and age group (*χ*^2^(1) = 0.22, *p* = 0.64). However, we found a similar pattern of how continuous age measures impact solution time: there was an effect that differed between children and adults (*χ*^2^(2) = 44.5, *p* = 2.2 * 10^−10^), with adults slowing down with age (on average taking an additional 1.16*s* per year, 95% CI=[0.87, 1.46], *χ*^2^(1) = 59.4, *p* = 1.3 * 10^−14^), and children becoming numerically but not statistically faster with age (on average taking 2.64*s* less per year, 95% CI=[-1.63, 6.90], *χ*^2^(1) = 1.47, *p* = 0.23).

**Figure 5.**
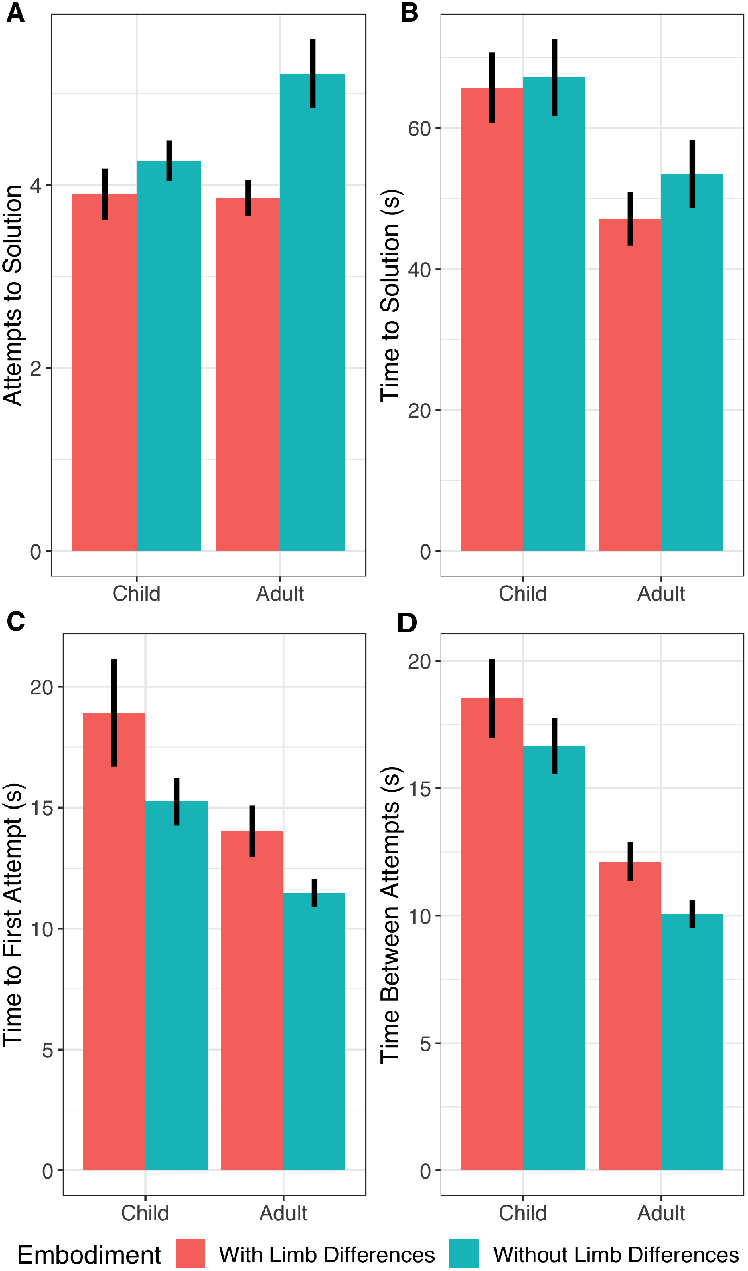
The efficiency of finding solutions measured by number of attempts (**A**), time to solution as measured in seconds (**B**), time to first attempt (**C**), and time between attempts (**D**). Means with standard errors are shown. Participants with and without limb differences did not reliably differ on time to solution, but participants with limb differences solved the levels in fewer attempts, and took more time until the first attempt and between attempts.

Thus we found that development and indeed aging cause noticeable changes in overall performance on the Virtual Tools game, but did not find that participants with or without limb differences are any better at the game. Nonetheless, participants might achieve similar overall levels of performance but do so in different ways. We therefore next considered whether participants with or without limb differences might demonstrate distinctions on the more detailed performance metrics specified in Allen et al. (2020).

### Both age and limb differences affect the way levels are solved

We investigated whether there was a difference in the number of attempts that participants with and without limb differences took to solve each level. Participants with limb differences (LD) took fewer attempts on average to come to a solution than the participants with no limb differences (NLD; 83.9% of the attempts, 95% CI=[72.2%, 97.6%]; *χ*^2^(1) = 5.19, *p* = 0.023; Figure 5A). Conversely, they took more time for each attempt, including thinking more before the first attempt (3.79*s* more, 95% CI=[1.99, 5.59]; *χ*^2^(1) = 17.0, *p* = 3.7 * 10^−5^; Figure 5C), and between all subsequent attempts (2.80*s* more on average, 95% CI=[1.51, 4.10]; *χ*^2^(1) = 17.9, *p* = 2.3 * 10^−5^; Figure 5D). Again, we found differences by age, with children taking fewer attempts (*χ*^2^(1) = 6.51, *p* = 0.011); more time to the first attempt (*χ*^2^(1) = 13.4, *p* = 0.00025); and more time between attempts (*χ*^2^(1) = 41.7, *p* =1.1 * 10^−10^), but no evidence for an interaction between age and limb differences for any of these measures (number of attempts: *χ*^2^(1) = 2.07, *p* = 0.15; time to first attempt: *χ*^2^(1) = 0.10, *p* = 0.76; time between attempts: *χ*^2^(1) = 0.16, *p* = 0.69).^1^

Together, these results suggest that individuals born with limb differences learn a different strategy for physical problem-solving: they learn to rely more on thinking about the problem and less on gathering information from their attempts.

### Analyzing types of attempts reveals group differences

We first investigated whether there were differences in the types of first attempts taken by participants with and without limb differences, using the methodology described in the Methods:Analysis section to classify whether the types of attempts made by participants with limb differences better matched those of other participants with limb differences than those without, and vice versa. If this measure is on average reliably above chance on a level, this suggests that the two groups are beginning their solution search in different ways. However, we did not find statistically reliable effects. For qualitative differences between groups see Figure 7, and Section S9 of the Supplemental Material for further details.

Using the same methodology, we asked whether children’s first attempt types could be differentiated from adults’. We found that across all levels, children’s and adults’ first attempt types can be differentiated (Fig 6B; note that in this cluster analysis comparing age groups, we cannot jointly model the effect of limb differences and age, so we report analyses separately for each group; NLD: *t*(84) = 4.91, *p* = 4.5 * 10^−6^; LD: *t*(57) = 3.05, *p* = 0.0035). Thus, children did not merely take more time and fewer attempts than adults – they also often chose different types of attempts to adults from the outset.

**Figure 6.**
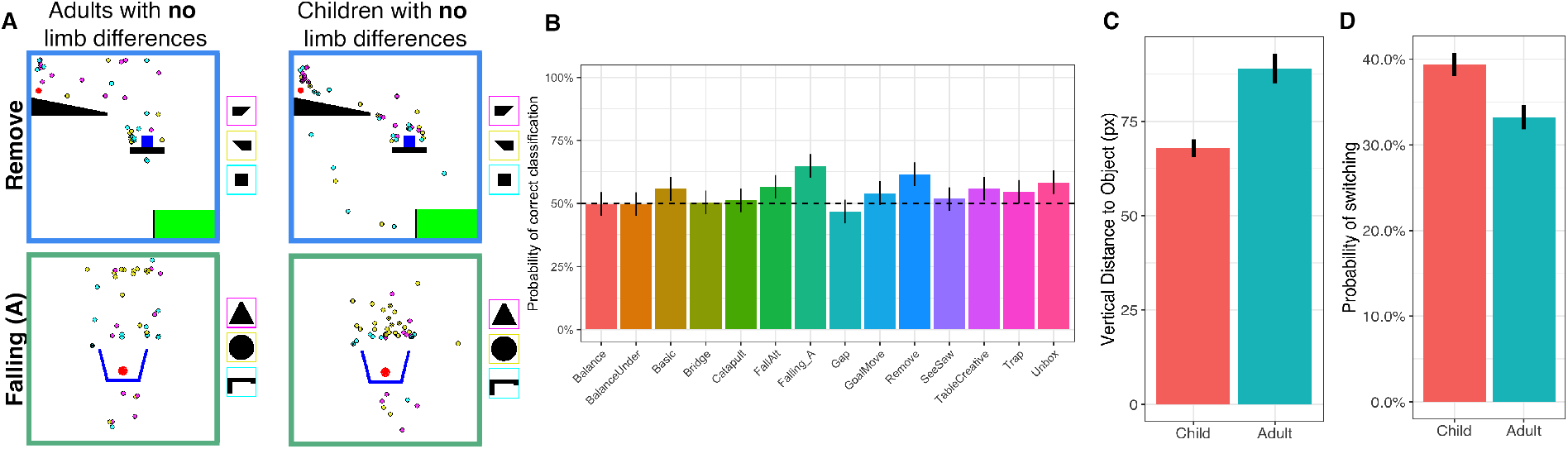
Comparing tool placements across children and adults born with two hands. Bar plots show means and standard errors. (**A**) Examples of first attempts for both adults and children without limb differences on two levels. Each point shows an individual participant’s attempt, with the position being where they placed the tool, and the color representing which tool they chose (tools shown in colored boxes to the right of each level). (**B**) We tested whether we could classify participants’ age group based on attempt type for each level. The dashed line represents 50% (chance). (**C**) The average vertical distance between the tool placement and the nearest moveable object. (**D**) The likelihood that children would “switch” strategies between attempts. Please see Methods:Analysis and Section S5 in the Supplemental Material for details on how strategy switching was measured.

In an exploratory analysis (Fig 6C and 7), we noticed that these differences specifically arose from the fact adults were more likely to place tools high above the objects they were trying to interact with, while children often dropped tools from closer. Quantifying this, we found that children tended to place the tools 19% closer to the objects that they intended to move or support than adults did (average vertical distance in children: 74px, adults: 91px; *χ*^2^(1) = 5.20, *p* = 0.023; Figure 6C). This was not simply due to a tendency to place tools nearer to objects in general, as there was no reliable difference in horizontal distance between tools and the nearest object (children: 31.3*px*, adults: 30.7*px*; *χ*^2^(1) = 0.65, *p* = 0.42). While speculative, it is possible that these vertical differences might be driven by differences in real-world experience between children and adults, perhaps because of additional experience with objects falling under gravity.

**Figure 7.**
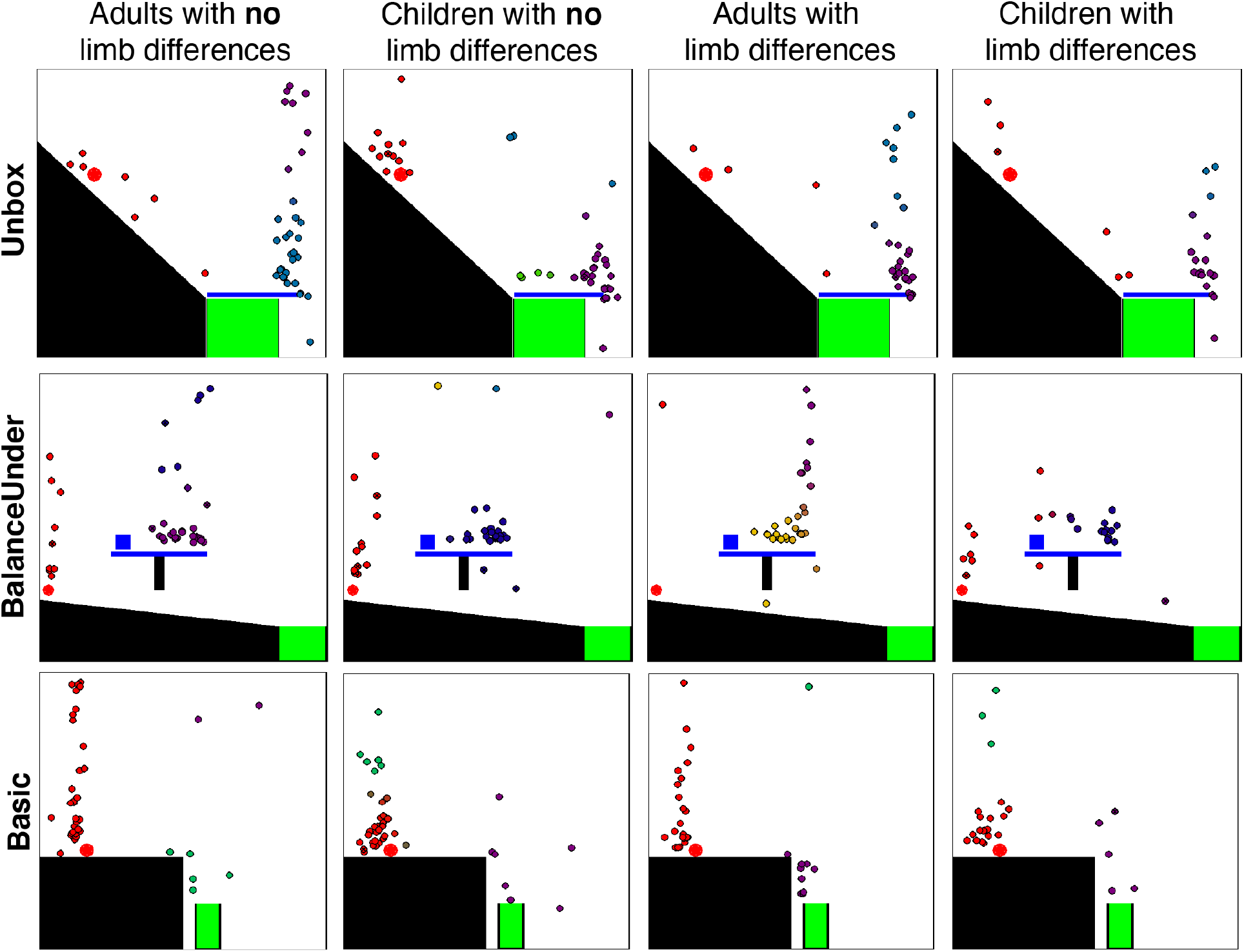
A comparison of tool placements on the first attempt for children and adults with and without limb differences. Each point represents an individual participant’s first attempt, with the position being where they placed the tool, and the color representing membership in “strategy” clusters determined by Dirichlet Process Mixture Modeling over tool positions. Points of the same color therefore represent participants who chose first attempts belonging to the same strategy.

We also investigated whether there was any difference in how children’s types of attempts changed through the course of a single level as compared to adults. Because children have been found to be more exploratory than adults (Gopnik, Sobel, Schulz, & Glymour, 2001), we tested for differences in exploration behavior: would children be more or less likely to stick with similar types of attempts to what they had just tried, or try something new? All participants’ attempts were used to form clusters to describe different “strategies” for using tools (see the Methods:Analysis section for details). We then assigned each attempt to one of these strategies (see Figure 7 for examples), and asked whether children or adults were more likely to switch strategies between attempts. This analysis demonstrated that children were more likely to try new strategies than adults (children: 39% strategy switches, adults: 33%; *χ*^2^(1) = 10.2, *p* = 0.0014), suggesting that their lower solution rates might be due to either increased exploration of inefficient strategies, or giving up on promising strategies early.

## Discussion

We used a virtual physics problem-solving game to study how growing up in a different body affects a high-level cognitive task unrelated to body or hand representations. We found minimal differences in the specific kinds of actions used by individuals with and without limb differences, suggesting that fundamental aspects of physical problem-solving are not dependent on similar kinds of manipulation experience. However, we also found that individuals with limb differences, regardless of age, spent more time thinking about virtual physical problems, and took fewer attempts to find solutions. While congenital limb difference is not directly associated with cognitive differences (see Section S2.1 in the Supplemental Material), growing up with a limb difference may cause cascading effects on many aspects of development, relating to different opportunities to interact with the environment. For example, children with limb differences may be unable to solve problems by imitating their parents or peers, or may face challenges in using tools designed for two hands. In either case, they might naturally come to appreciate the value of thinking more about solutions to physical problems before acting, and over time this could grow into a general strategy for interacting with the world. What is striking is that this learned strategy extends to a task in which action possibilities are equated across groups – indeed, individuals without limb differences were not slower to control the cursor in our task (see also Maimon-Mor et al., 2021).

Such cognitive strategies may be learned through experience, similarly to studies of motor learning in adults (Huang, Kram, & Ahmed, 2012). The difference here is that the target of learning is not the motor plan itself, but *when to deploy* those motor plans. While people’s information sampling has been shown to be sensitive to the costs of obtaining that information (Juni, Gureckis, & Maloney, 2016; Jones et al., 2019) and motor cost manipulations have been shown to affect the efficiency of motor reaching actions (Summerside, Shadmehr, & Ahmed, 2018), it has not previously been shown that motor differences directly affect the cognitive strategies that people employ.

Our findings also bridge two different approaches to understanding human tool use. Tool use is theorized to be supported by specific sensorimotor knowledge of tool manipulation under the “manipulation-based” (embodied cognition) approach (Buxbaum & Kalénine, 2010; Gonzalez Rothi, Ochipa, & Heilman, 1991; van Elk, van Schie, & Bekkering, 2014), or generic physical knowledge under the “reasoning-based” approach (Allen et al., 2020; Osiurak & Badets, 2016). These theories have produced suggestions that there are distinct cognitive systems supporting different kinds of tool knowledge (Orban & Caruana, 2014; Goldenberg & Spatt, 2009). However, our results suggest a connection between the two systems: by its virtual nature and novel objects, the Virtual Tools game must rely on reasoning-based systems for tool use, yet we find that manipulation capabilities affect this reasoning. Thus we suggest that the development of the reasoning-based system is grounded in the embodied way that we interact with the world.

Developmentally, our results extend existing knowledge about children’s problem-solving and tool use. While even preschoolers (Gopnik et al., 2001) or infants (Baillargeon, Li, Gertner, & Wu, 2011) understand cause and effect, our task involves *reasoning* about the effects of the virtual tools on their environment. Children can use and select known tools by 2-3 years old (Keen, 2011), but do not reliably innovate new tools until 8-9 years (Beck et al., 2011), perhaps because innovation requires increased cognitive demands, including creativity, attentional control, inhibition, and planning (Rawlings & Legare, 2020). Children’s performance in our game is likely driven by these same skills: like complex tool-use or innovation, children must strike a balance between exploration and exploitation in a large solution space (Gopnik, 2020). Specifically, children must avoid overcommitting to a single suboptimal approach, but also must not explore the space too much, lest they lose track of promising solutions. Children at this age can avoid perservation (Rawlings & Legare, 2020; Cutting, Apperly, & Beck, 2011), and our results suggest over-exploration is the more likely trap, as children switched attempt types more often than adults. It may be that the central difficulty with tool innovation and other physical problem-solving tasks at this age is the need to search through large solution spaces, where children’s propensity for exploration (Oudeyer & Smith, 2016; Gopnik, 2020) may come at the cost of short-term gains in solution-finding.

Our findings suggest new interactions between embodiment, development, and cognitive strategies and raise important questions for future work. The present study focused on a single task – the Virtual Tools game – because it is similar to manipulation tasks while still not requiring manipulation directly. Thus we expected cognitive strategies learned from lifelong differences in action costs and possibilities to carry over into this task, while still providing an equal playing field for all our participants.

One might ask how broadly these differences in cognitive strategies would be expected to generalize, e.g. across domains and environments. Some previous work indirectly supports the notion of a broad effect across motor-related tasks. For example, individuals with a limb difference responded more slowly in motor planning (Philip et al., 2015) and hand laterality judgements (Maimon-Mor et al., 2020). These findings were previously interpreted as a consequence of having fewer available resources to accumulate evidence for tasks related to judgements about hands. However, in light of the present findings, and because the previous observed effects were also found for the intact hand, these results can be interpreted as a *generally* greater reliance on planning before acting for this population. Yet, these tasks all contain a visuospatial and motor component, so it is still unclear how broad this effect is, e.g., whether it transfers to abstract logical reasoning tasks or social interactions. It is also difficult to determine, based on our single study, whether the group differences we observed are directly or indirectly caused by different embodied experience considering the many developmental differences individuals growing up with a different body will experience. For example, life experiences for individuals born with different bodies may cause them to be generally more cautious, contemplative, or creative.

Thus, the current study provides a starting point for further investigation of how we learn to deploy our cognitive resources based on our embodied experiences. Being born with a different body does not change the fundamental ways in which people try to act on the world, but it can change the strategies they learn in order to plan and act efficiently in their environments.

## Acknowledgements

We thank Mathew Kollamkulam for collecting the data from the adult samples, Opcare, Limbo and Reach for assistance with participant recruitment, and all participating families and volunteers. The study was supported by a Wellcome Trust Senior Research Fellowship (215575/Z/19/Z), awarded to TRM. KRA, KAS, and JBT were supported by National Science Foundation Science Technology Center Award CCF-1231216; Office of Naval Research Multidisciplinary University Research Initiative (ONR MURI) N00014-13-1-0333; and research grants from ONR, Honda, and Mitsubishi Electric. For the purpose of Open Access, the author has applied a CC BY public copyright licence to any Author Accepted Manuscript (AAM) version arising from this submission.

## Supplemental Material

### S1 Additional experimental details

#### S1.1 Virtual Tools Game

**Figure S1.**
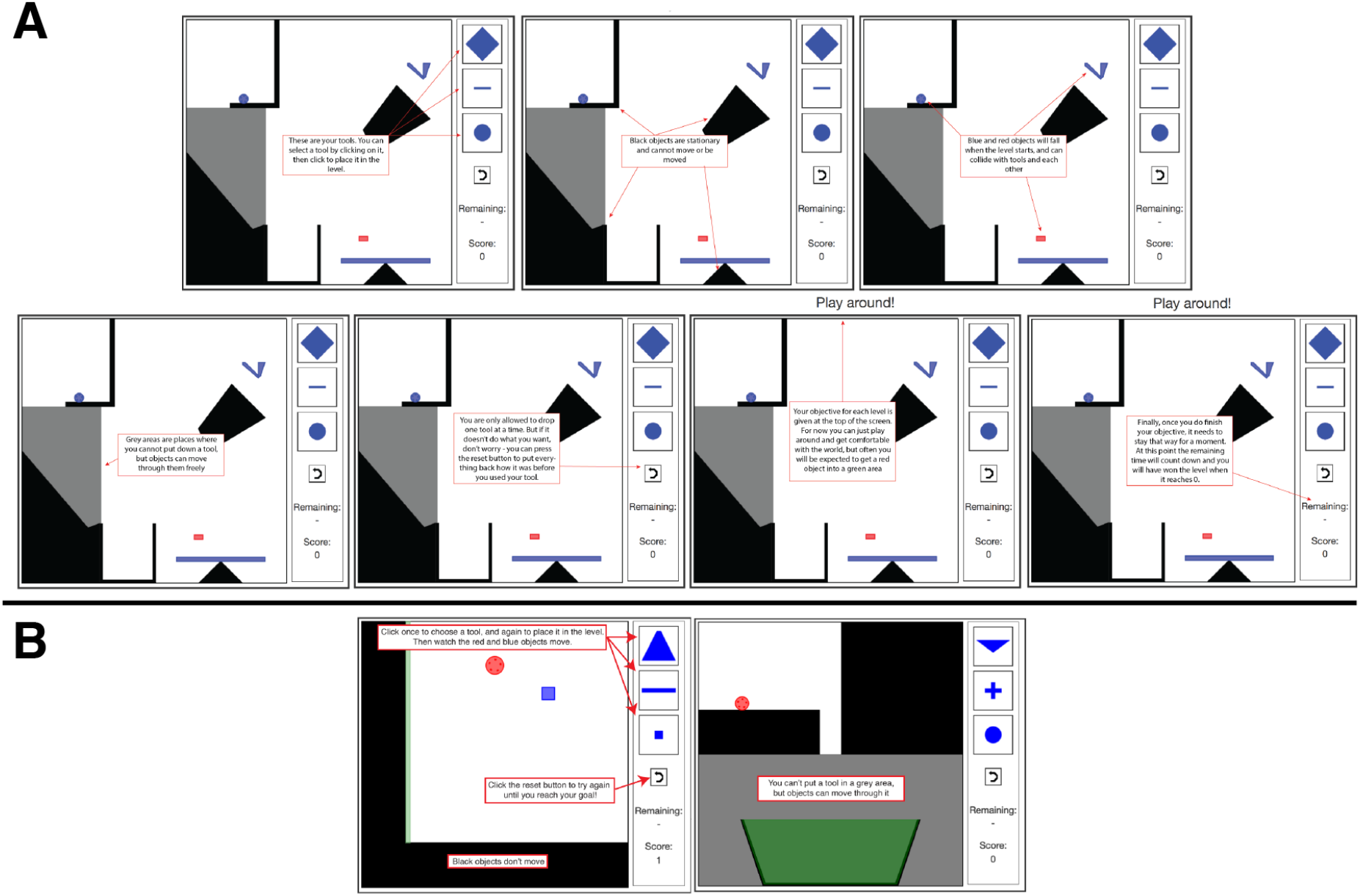
Instruction screens provided to **A.** adults and **B.** children.

The instructions provided to children and adults for the Virtual Tools game are provided in Figure S1. Introductory levels which must be solved (by both children and adults) but were not analyzed were the levels shown in the childrens’ instructions.

#### S1.2 Motor pre-test

The motor pre-test ensured that participants with and without limb differences had equal abilities to interact with the computer system, and therefore should be equally able to control the Virtual Tools game. The motor-test is shown in Figure S2. Participants were required to click a central star before clicking a colored circle in the periphery (either 150 or 250 px from the center of a 600×600px screen). They completed 10 rounds of this procedure.

We presented median motor RT (our motor covariate) as Figure 3 in the main text, but see Figure S4 for median motor error (in pixels).

**Figure S2.**
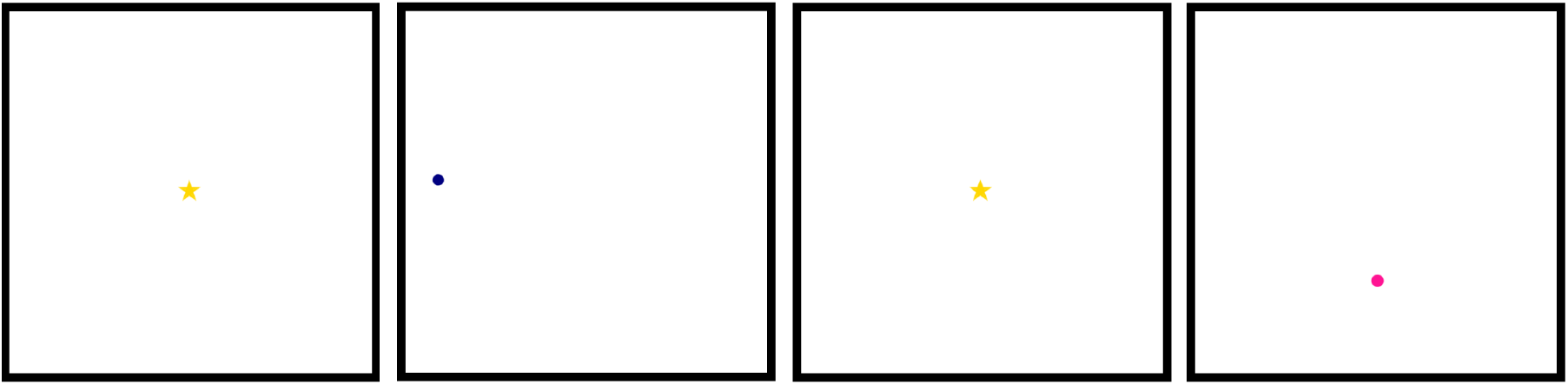
An example of two rounds of the motor pre-test. Participants clicked first the star, then a circle in the periphery, as quickly as possible.

### S2 Participant demographics

#### S2.1 Education levels

Adults were matched on education level, and we tested for similarities in cognitive capabilities by performing IQ tests on a subset of the children from both the LD and NLD groups. These tests were provided in a separate session from the main experiment. We assessed both Raven’s matrices measures of spatial IQ, and BPVS as a measure of verbal IQ. 20/25 Child-LD and 34/43 Child-NLD participants were tested. All scores were within normal range (lowest for Ravens 85, highest 135, lowest for BPVS 75, highest 134. Ravens means: Child-LD: 113, Child-NLD: 111 for Ravens. BPVS means Child-LD: 105, Child-NLD: 108), and 2-sided unpaired t-tests show no evidence for differences between these populations (Ravens: *t*(40) = 0.5, *p* = 0.6, BPVS: *t*(50) = 0.8, *p* = 0.4).

#### S2.2 Demographic and interface information

Please see Table S1 for information on participant handedness as well as how they interfaced with the computer.

**Table S1.** Demographics summaries for participants. LD: limb differences; NLD: no limb differences

| Group | Mean Age (SD) | Handedness |  |  | Input Device |  |  |
| --- | --- | --- | --- | --- | --- | --- | --- |
|  |  | Left | Right | Ambi. | Mouse | Touchpad | Other |
| Adult-LD | 40.7 (15.5) | 0.45 | 0.55 | 0 | 0.61 | 0.36 | 0.03 |
| Adult-NLD | 41.2 (15.2) | 0.08 | 0.92 | 0 | 0.55 | 0.45 | 0 |
| Child-LD | 7.91 (1.84) | 0.36 | 0.64 | 0 | 0.32 | 0.68 | 0 |
| Child-NLD | 7.89 (1.73) | 0.07 | 0.89 | 0.04 | 0.50 | 0.50 | 0 |

**Figure S3.**
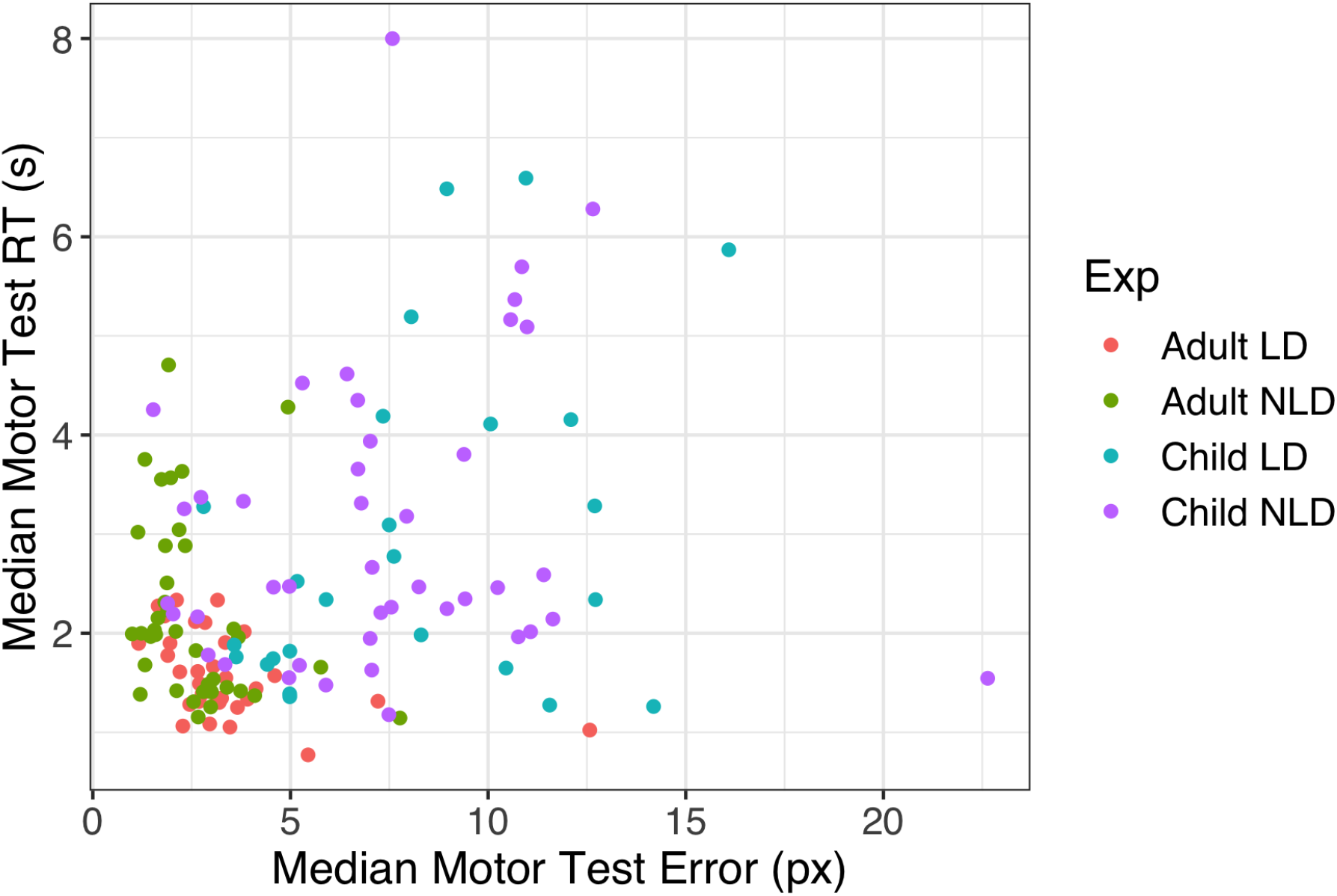
Comparison of Median Motor test reaction time with median motor test error for each participant across groups. LD = participant with limb differences, NLD = participant with no limb differences. Correlation is reasonably high, *r* = 0.34, so we use only the median motor test error as a covariate throughout analyses.

#### S2.3 Specific limb differences

For details on specific limb differences in the children we tested, please refer to Table S3. Adult participants had less variability in their limb differences. Of the 33 adult participants, one had a bilateral limb difference (with a missing right arm at the shoulder, and left arm at the elbow), and 32 had unilateral limb differences, including:

- 3 transhumeral
- 3 at the elbow
- 18 transradial (one participant has a small residual digit)
- 8 at wrist (one participant has 5 small intact digits; one has a short thumb and small, non-jointed digit; one has two residual digits)

**Table S2.** Details of adults with a limb difference. Pincer: defined as the ability to grasp a small object between the thumb and index finger; Dominant limb: R=right, L=left. For prosthesis usage, 1 indicates daily usage > 8 hours per day, 2 indicates daily usage 4-8 hours per day, 3 indicates daily less than 4 hours per day, 5 indicates rare usage.

| ID | Limb Difference |  |  |  | Dominant Limb | Prosthesis |  |
| --- | --- | --- | --- | --- | --- | --- | --- |
|  | Left | Pincer | Right | Pincer |  | Fitted | Use |
| 1 | Intact | Yes | Absent below elbow | No | L | Yes | 1 |
| 2 | Absent above elbow | No | Intact | Yes | R | No | - |
| 3 | Absent above elbow | No | Intact | Yes | R | Yes | 2 |
| 4 | Absent below elbow | No | Intact | Yes | R | Yes | 2 |
| 5 | Absent below elbow | No | Intact | Yes | R | Yes | 2 |
| 6 | Absent below elbow | No | Intact | Yes | R | Yes | 1 |
| 7 | Intact | Yes | Absent below elbow | No | L | Yes | 1 |
| 8 | Intact | Yes | Absent below elbow | No | L | No | - |
| 9 | Absent below elbow | No | Intact | Yes | R | Yes | 2 |
| 10 | Absent below wrist | No | Intact | Yes | R |  |  |
| 11 | Absent below elbow | No | Intact | Yes | R | Yes | 2 |
| 12 | Absent below elbow | No | Intact | Yes | R | Yes | 1 |
| 13 | Intact | Yes | Absent below elbow | No | L | Yes | 2 |
| 14 | Intact | Yes | Absent below elbow | No | L | Yes | 2 |
| 15 | Absent below elbow | No | Intact | Yes | R | Yes | 1 |
| 16 | Intact | Yes | Absent below elbow | No | L |  |  |
| 17 | Absent below elbow | No | Intact | Yes | R | Yes | 1 |
| 18 | Absent below elbow | No | Intact | Yes | R | Yes | 3 |
| 19 | Absent below wrist | No | Intact | Yes | R | No | - |
| 20 | Absent below elbow | No | Intact | Yes | R | Yes | 1 |
| 21 | Intact | Yes | Absent below elbow | No | L | No | - |
| 22 | Intact | Yes | Absent below elbow | No | L | Yes | 1 |
| 23 | Intact | Yes | Absent below wrist | No | L | Yes | 2 |
| 24 | Partial hand, 1 digit | No | Intact | Yes | R | Yes | 5 |
| 25 | Intact | Yes | Absent above elbow | No | L | Yes | 5 |
| 26 | Absent below elbow | No | Absent below shoulder | No | L | No | - |
| 27 | Intact | Yes | 5 small digits | Yes | L | No | - |
| 28 | Intact | Yes | Short thumb, 1 non-jointed digit | Yes | L | No | - |
| 29 | Partial hand, 2 digits | Yes | Intact | Yes | R |  |  |
| 30 | Absent below elbow | No | Intact | Yes | R | No | - |
| 31 | Intact | Yes | Absent below wrist | No | L | Yes | 5 |
| 32 | Intact | Yes | Absent below wrist | No | L | No | - |
| 33 | Absent below elbow | No | Intact | Yes | R | Yes | 2 |

**Table S3.** Details of children with a limb difference. Pincer: defined as the ability to grasp a small object between the thumb and index finger; Dominant limb: R=right, L=left; Prosthesis use is measured as 1: use everyday, 2: use 5 days a week, 3: use weekly, 4: use monthly, 5: use a few times a year.

| ID | Limb Difference |  |  |  | Dominant Limb | Prosthesis |  |
| --- | --- | --- | --- | --- | --- | --- | --- |
|  | Left | Pincer | Right | Pincer |  | Fitted | Use |
| 1 | Partial hand, 1 digit | No | Partial hand, 1 digit | No | R | No | - |
| 2 | 5 digits, limited function | Limited | Intact | Yes | R | No | - |
| 3 | Partial hand, 1 digit | No | Partial hand, 3 digits | Yes | R | No | - |
| 4 | Absent below elbow | No | Intact | Yes | R | Yes | 5 |
| 5 | Absent below elbow | No | Intact | Yes | R | Yes | 6 |
| 6 | Intact | Yes | Partial hand, 2 digits | Yes | L | No | - |
| 7 | Partial hand, 3 digits | No | Partial hand, 4 digits | No | R | No | - |
| 8 | Partial hand, no digits | No | Intact | Yes | R | Yes | 2 |
| 9 | Intact | Yes | Absent below elbow | No | L | Yes | - |
| 10 | Absent below elbow | No | Intact | Yes | R | No | - |
| 11 | Absent below elbow | No | Intact | Yes | R | Yes | 3 |
| 12 | Intact | Yes | Partial hand, no digits | No | L | No | - |
| 13 | Partial hand, 3 digits | Limited | Partial hand, 3 digits | Yes | R | No | - |
| 14 | 5 digits, non-functional | No | Intact | Yes | R | No | - |
| 15 | Intact | Yes | Partial wrist, no digits | No | L | Yes | - |
| 16 | Intact | Yes | Partial hand, 2 digits | Yes | L | No | - |
| 17 | 4 digits, non-functional | No | 4 digits, non-functional | No | R | No | - |
| 18 | Absent below elbow | No | Intact | Yes | R | Yes | 5 |
| 19 | Intact | Yes | Absent below elbow | No | L | No | - |
| 20 | Absent above elbow | No | Absent above elbow | No | L | No | - |
| 21 | Intact | Yes | Absent above elbow | No | L | No | - |
| 22 | Absent below wrist | No | Intact | Yes | R | Yes | 2 |
| 23 | Absent below elbow | No | Intact | Yes | R | Yes | 3 |
| 24 | Intact | Yes | Partial hand, 4 digits | Yes | L | No | - |
| 25 | Partial hand, 2 digits | No | Intact | Yes | R | No | - |

**Figure S4.**
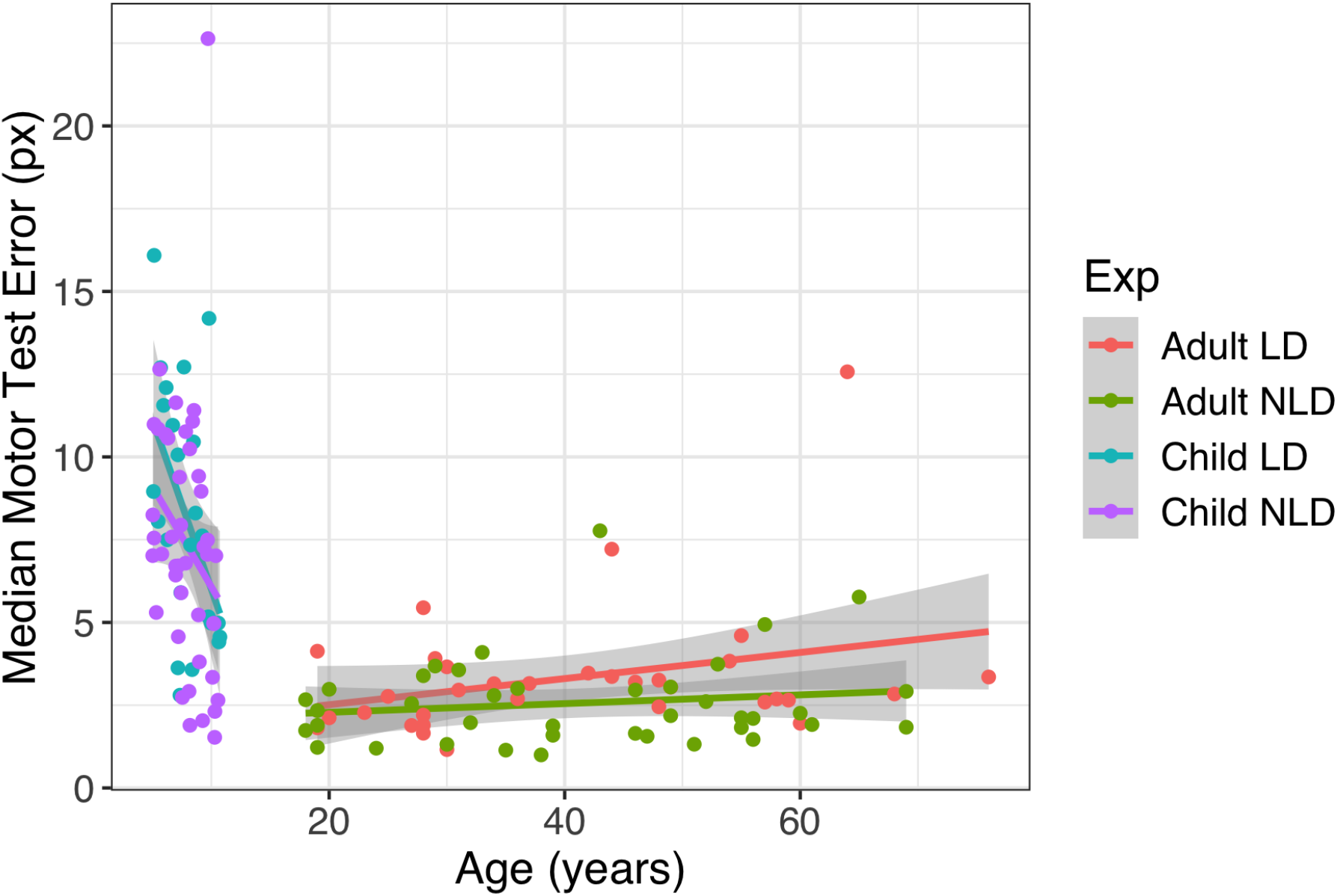
Median motor test error in pixels as a function of age. Children improve significantly as they age, while adults’ performance is unaffected. No interaction was found between age and embodiment group ((*F*(1, 134) = 0.001, *p* = 0.98)).

### S3 Exclusions

We excluded two adults with limb differences from analysis as they were amputees rather than having congenital limb differences. We also excluded results from two two-handed children: one due to a data recording error, and one who provided unreliable motor test data due to continuously clicking rather than attempting the task.

Adult participants were given 15 different levels to solve: the 14 shown in Fig 2C, and one additional level: Spiky (see Allen et al., 2020). Because of the low solution rate of adults on this level (18%), we were concerned that it might frustrate children and cause attrition, so only had children play the other 14 levels. To keep the groups matched, we therefore removed the Spiky level from analysis.

### S4 Analysis details

We modeled all statistical analyses as (generalized) linear mixed effect models using the ‘lme4’ package in R (Bates, Mächler, Bolker, & Walker, 2015). We treated accuracy as a binomial response, time measures as having Gaussian error, and attempts as a Poisson process (using the number of non-solution attempts as the dependent variable so that we could observe zero-attempt outcomes). In all models, we assumed random intercepts for participants and levels.

Additionally, we prespecified two covariates for all of our analyses. We used the median motor test response time as a covariate, as we had hypothesized that motor facility might cause better performance. We selected response time instead of error because the two measures were correlated (*r* = 0.34), and in pilot analyses we found that adding a second motor measure explained very small amounts of additional variance in performance over just a single measure. Finally, because we assumed that there would be large effects of age on performance, for all analyses testing the difference between participants with and without limb differences, we included age as a covariate, allowing its effects on performance to differ for children and adults (treated as an age by child/adult interaction).

### S5 Attempt clustering procedure

To measure exploratory behavior, in the Results we looked at how likely participants were to “switch” strategies between attempts. Since strategies are hard to quantify, we used a nonparametric clustering package implemented in Python’s SciPy package (Virtanen et al., 2020) to discover strategy clusters from participant data. For each different level, we aggregated all attempts across all participants, taking the [*x, y*] spatial positions of each attempt as variables, but ignoring the specific choice of tool. We applied clusters separately for each participant group (children vs. adults, limb differences vs. no limb differences), but aggregated data across all participants from each group. Applying the Dirichlet Process mixture modeling package then gave us clusters for each level and group, as well as the probabilities that each attempt belonged to each cluster. We defined a “switch” as occurring if, for consecutive attempts, the cluster assigned as the highest likelihood were different for both of those attempts. Examples of discovered clusters for the data presented in Figure 7 are shown in Figure S5.

### S6 Analysis excluding covariates

Because we expected that age and motor capabilities would have strong effects on task performance, we used both continuous age measures and performance on the motor pre-test as covariates when testing for the effect of limb differences. However, because this experiment does not involve random assignment of participants to this dependent variable, we can and do find differences (albeit small) in these covariates across groups with and without limb differences. We therefore have additionally analyzed all primary measures *without* covariates to determine the effect of this analysis choice, and find qualitatively identical results, as reported in detail below. While there is a difference between children and adults on overall solution rate (*χ*^2^(1) = 12.0, *p* = 0.00053) and time to solve (*χ*^2^(1) = 13.9, *p* = 0.00019), there remains no effect of limb differences on either (solution rate: *χ*^2^(1) = 1.3, *p* = 0.256, solution time: *χ*^2^(1) = 0.06, *p* = 0.803), nor is there an interaction between limb differences and age group (solution rate: *χ*^2^(1) = 0.00, *p* = 0.944, time to solve: *χ*^2^(1) = 0.46, *p* = 0.498).

We still find that participants with limb differences take fewer attempts (*χ*^2^(1) = 5.88, *p* = 0.015) but more time thinking before taking their first attempt (*χ*^2^(1) = 7.00, *p* = 0.0081) and between subsequent attempts (*χ*^2^(1) = 4.68, *p* = 0.031, though no interaction between limb differences and age group (number of attempts: *χ*^2^(1) = 2.21, *p* = 0.137, time to first attempt: *χ*^2^(1) = 0.286, *p* = 0.593, time between attempts: *χ*^2^(1) = 0.01, *p* = 0.915). Similar to the analysis with covariates, adults take less time to the first attempt (*χ*^2^(1) = 13.09, *p* = 0.0003) and between attempts (*χ*^2^(1) = 42.9, *p* = 5.6 * 10^−11^).

### S7 Controlling for possible moderators

In order to validate our choice of analysis, we explored whether any of a set of moderators might have driven any of the results we report in the paper. These moderators were not expected to have an impact on the results, but were tested in exploratory analysis to check that assumption. We specifically test (1) participant gender, (2) the type of device participants used to control the game (mouse or touchpad), (3) whether participants were dominantly left- or right-handed, and (4) for the DL participants, what kind of prostheses they used. While we find possible impacts of these variables on overall performance (as detailed in the next paragraph), in no cases do we find any interactions between these variables and embodiment, suggesting that they should not impact the main results of the paper.

Gender had a main effect on solution rate (*χ*^2^(1) = 9.88, *p* = 0.0017), with males slightly outperforming females (83% vs 77%), but no interaction with embodiment (*χ*^2^(1) = 0.53, *p* = 0.47). We also found a small effect of gender on the first attempt time (males: 13.8*s*, females: 15.2*s*, *χ*^2^(1) = 5.56, *p* = 0.018), but again no interaction with embodiment (*χ*^2^(1) = 0.75, *p* = 0.39). There were no other main effects of gender or interactions between gender and embodiment for any other performance metrics. Device type had a main effect on time to solution (*χ*^2^(1) = 7.96, *p* = 0.0048), with participants using a mouse solving the levels slightly faster than participants using a touchpad (53.2*s* vs 63.8*s*). However, we found no interaction between device and embodiment (*χ*^2^(1) = 0.47, *p* = 0.49), nor did we find any other main effects of input device type. We found no effect of hand laterality on any of our dependent variables. We similarly find no main or interaction effects for prosthesis use on any of our performance measures (all *ps >* 0.12).

### S8 Analysis of all levels

In the main body of the text, we studied the impact of age and embodiment on how participants solved *successful* levels in order to avoid measuring effects of persistence or motivation. In order to test the effect of this analysis choice on our results, we additionally ran our main analysis including all levels. We find that in general, there is little difference between the success-conditioned and all-levels analyses, though one statistical test (overall number of attempts by embodiment) crosses from statistically significant to barely not statistically significant. Based on exploratory analysis, this appears to be driven by differences in perseveration between groups, with participants with limb differences persisting marginally more than those without. We discuss this difference and report all statistics below for transparency.

#### S8.1 Number of total attempts

While we found that participants with limb differences used fewer attempts to get to a solution than participants without, the effect of embodiment on number of total attempts is only marginally significant (participants with limb differences taking 88.1% of attempts of those without on average, 95% CI=[86.1%, 102.0%]; *χ*^2^(1) = 2.92, *p* = 0.088).

We ask whether any differences in this analysis might be a function of differences in motivation, and so measure persistence as the ratio of total attempts (regardless of success) to the number of attempts taken on successful levels. We find that by this definition, participants with limb differences were numerically more persistent (adults: 145%, children: 147%) than participants with no limb differences (adults: 132%, children: 136%), though this difference was only marginally significant (*F*(1, 135) = 3.80, *p* = 0.053). Because of the marginal significance, we do not make strong claims about the persistence of LD vs. NLD participants, but this numerical difference is likely what drives the reduction in the measured effect of embodiment on total attempts.

#### S8.2 Other results

*Time to solution*: similar to the success-conditioned analysis, we find that children take more time than adults (*χ*^2^(1) = 37.8, *p* = 7.8 * 10^−10^), that there is no reliable effect of embodiment (*χ*^2^(1) = 0.37, *p* = 0.55), and that there is a difference by continuous age (*χ*^2^(2) = 43.8, *p* = 3.1 * 10^−10^). *Time to first attempt*: just as with the success-conditioned results, we find that participants with limb differences took more time than participants with no limb differences until their first attempt (4.12*s* more, 95%CI = [2.20, 6.03]; *χ*^2^(1) = 17.7, *p* = 2.6 * 10^−5^), and that this differs by age (*χ*^2^(1) = 12.4, *p* = 0.00043), but no evidence of an interaction between age and embodiment (*χ*^2^(1) = 0.32, *p* = 0.57). *Time between attempts*: similar to the success-conditioned analyses, we find that participants with limb differences spend more time thinking than participants with no limb differences between attempts (2.33*s* more, 95%CI = [1.04, 3.63]; *χ*^2^(1) = 12.5, *p* = 0.00041), and that this differs by age (*χ*^2^(1) = 43.5, *p* = 4.2 * 10^−11^), but no evidence of an interaction between age and embodiment (*χ*^2^(1) = 0.04, *p* = 0.84).

**Table S4.**
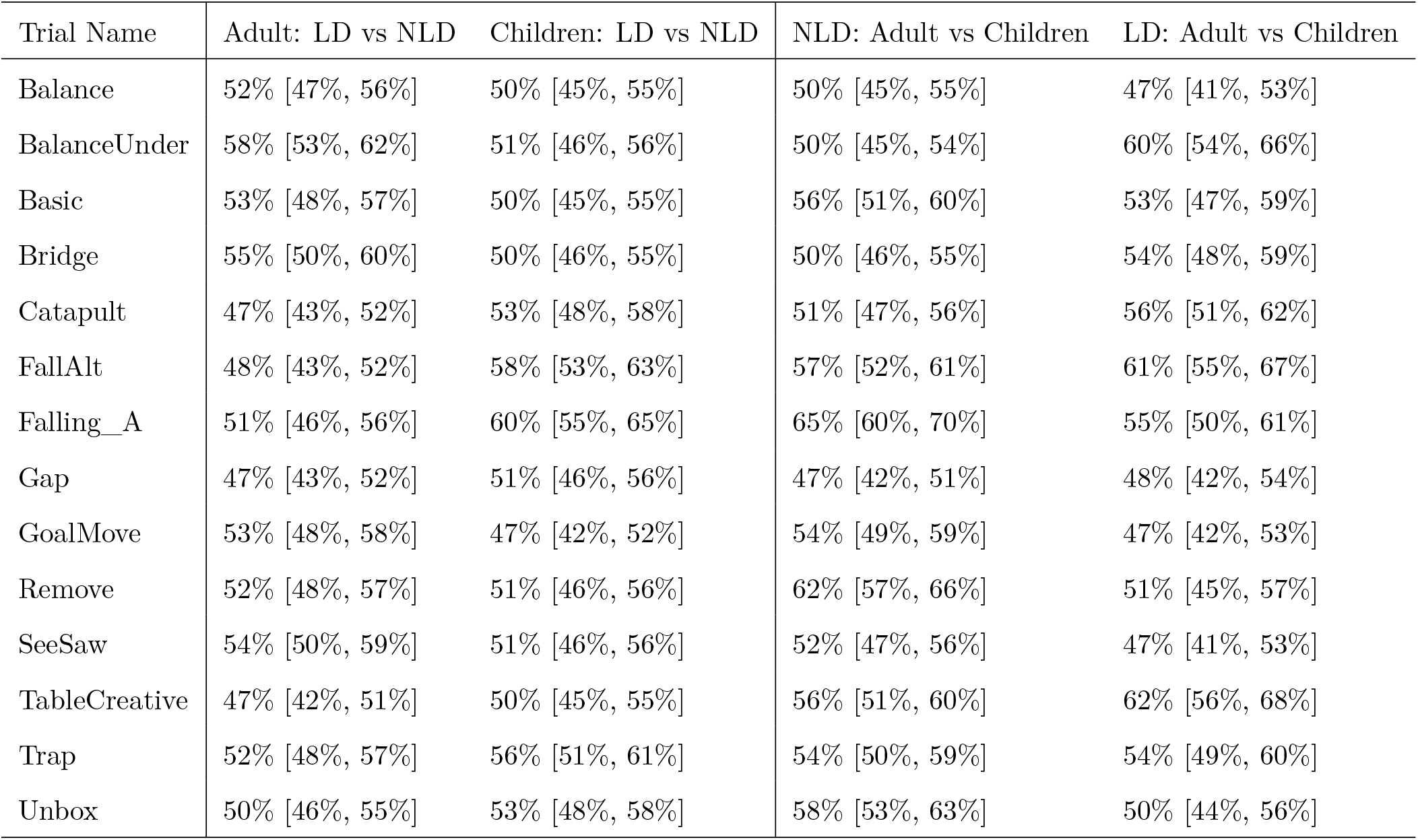
Classification of first attempts between different groups. Numbers in brackets represent bootstrapped 95% confidence intervals on classification percentages.

| Trial Name | Adult: LD vs NLD | Children: LD vs NLD | NLD: Adult vs Children | LD: Adult vs Children |
| --- | --- | --- | --- | --- |
| Balance | 52% [47%, 56%] | 50% [45%, 55%] | 50% [45%, 55%] | 47% [41%, 53%] |
| BalanceUnder | 58% [53%, 62%] | 51% [46%, 56%] | 50% [45%, 54%] | 60% [54%, 66%] |
| Basic | 53% [48%, 57%] | 50% [45%, 55%] | 56% [51%, 60%] | 53% [47%, 59%] |
| Bridge | 55% [50%, 60%] | 50% [46%, 55%] | 50% [46%, 55%] | 54% [48%, 59%] |
| Catapult | 47% [43%, 52%] | 53% [48%, 58%] | 51% [47%, 56%] | 56% [51%, 62%] |
| FallAlt | 48% [43%, 52%] | 58% [53%, 63%] | 57% [52%, 61%] | 61% [55%, 67%] |
| Falling_A | 51% [46%, 56%] | 60% [55%, 65%] | 65% [60%, 70%] | 55% [50%, 61%] |
| Gap | 47% [43%, 52%] | 51% [46%, 56%] | 47% [42%, 51%] | 48% [42%, 54%] |
| GoalMove | 53% [48%, 58%] | 47% [42%, 52%] | 54% [49%, 59%] | 47% [42%, 53%] |
| Remove | 52% [48%, 57%] | 51% [46%, 56%] | 62% [57%, 66%] | 51% [45%, 57%] |
| SeeSaw | 54% [50%, 59%] | 51% [46%, 56%] | 52% [47%, 56%] | 47% [41%, 53%] |
| TableCreative | 47% [42%, 51%] | 50% [45%, 55%] | 56% [51%, 60%] | 62% [56%, 68%] |
| Trap | 52% [48%, 57%] | 56% [51%, 61%] | 54% [50%, 59%] | 54% [49%, 60%] |
| Unbox | 50% [46%, 55%] | 53% [48%, 58%] | 58% [53%, 63%] | 50% [44%, 56%] |

### S9 Analysis of differences between LD and NLD attempt types

Overall, we found a trend towards being able to classify participants with and without limb differences (mean adult classification accuracy: 51.4%, 95% CI=[49.9, 52.9], *t*(71) = 1.91, *p* = 0.060; mean child classification accuracy: 52.3%, 95% CI=[49.5%, 55.1%, *t*(70) = 1.62, *p* = 0.109), but found only one level where the confidence interval on the estimated classification probability exceeded chance for both children and adults (see Table S4): BalanceUnder. This level requires preventing objects from falling by placing a tool as a counterweight to another object, which might be a strategy the participants with limb differences used more often in daily life, as they are have to rely on their residual arm (which is shorter than their intact arm) when manipulating objects bimanually. However, given that the estimated classification probability is not far from chance and that we cannot reliably classify attempts in other levels that rely on balancing, future work would need to investigate particular differences in strategies learned from interaction with the environment.

**Figure S5.**
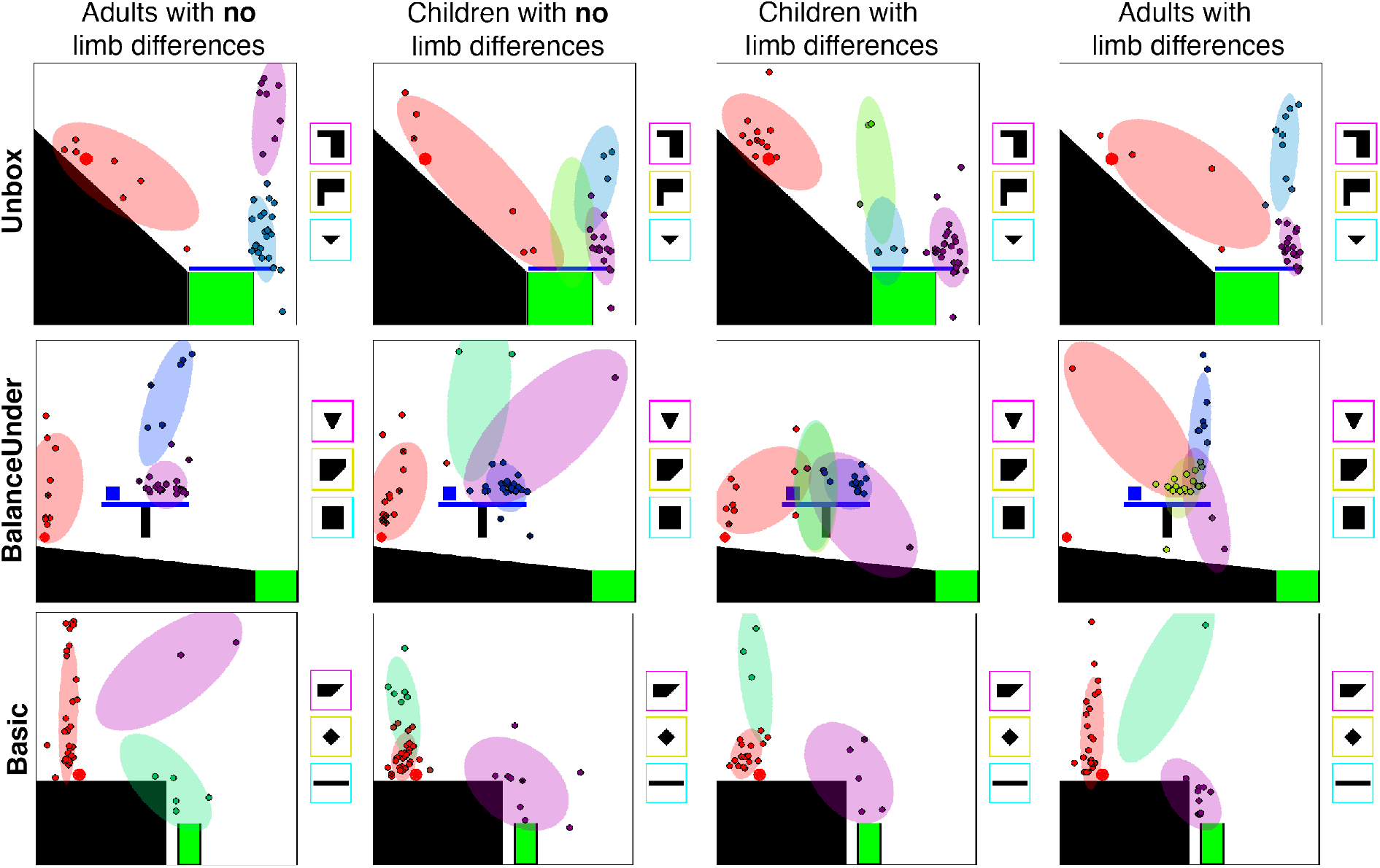
Examples of discovered strategy clusters (shown as semi-transparent ovals) with participants’ first attempts overlaid as colored points. The color of each point represents the highest likelihood cluster for that attempt.

## Footnotes

1 When testing for the effects of limb differences in children and adults separately, we found reliable differences in attempt timing in both children (first attempt: 4.13*s*, 95% CI=[0.96, 7.30], *χ*^2^(1) = 6.52, *p =* 0.011; between attempts: 2.57*s*, 95% CI=[0.31, 4.84], *χ*^2^(1) = 4.96, *p* = 0.026) and adults (first attempt: 3.29*s*, 95% CI=[1.32, 5.26], *χ*^2^(1) = 10.7, *p* = 0.0011; between attempts: 2.63*s*, 95% CI=[1.26, 4.01], *χ*^2^(1) = 14.2, *p* = 0.00017). However, while we found that adults with limb differences use fewer attempts than those without (75.1% of the attempts, 95% CI=[59.1%, 95.4%], *χ*^2^(1) = 5.49, *p* = 0.019), children with limb differences took numerically fewer attempts than those without, but this does not reach statistical significance (94.9% of the attempts, 95% CI=[78.2%, 115.0%], *χ*^2^(1) = 0.29, *p* = 0.59). Thus while we can claim that participants with no limb differences overall took more attempts, we did not have enough evidence to discriminate whether this is because these differences are consistent across age groups, or whether the distinction grows through development.

